# Emerging Evidence of Chromosome Folding by Loop Extrusion

**DOI:** 10.1101/264648

**Authors:** Geoffrey Fudenberg, Nezar Abdennur, Maxim Imakaev, Anton Goloborodko, Leonid A. Mirny

**Affiliations:** Gladstone Institutes, University of California, San Francisco, California, USA.; Computational and Systems Biology Program, Massachusetts Institute of Technology, Cambridge, Massachusetts, USA.; Institute for Medical Engineering and Science (IMES), Massachusetts Institute of Technology, Cambridge, Massachusetts, USA.; Department of Physics, Massachusetts Institute of Technology, Cambridge, Massachusetts, USA.

**Keywords:** chromatin, genome architecture, loop extrusion, Hi-C, polymer physics, CTCF, cohesin

## Abstract

Chromosome organization poses a remarkable physical problem with many biological consequences: how can molecular interactions between proteins at the nanometer scale organize micron-long chromatinized DNA molecules, insulating or facilitating interactions between specific genomic elements? The mechanism of active loop extrusion holds great promise for explaining interphase and mitotic chromosome folding, yet remains difficult to assay directly. We discuss predictions from our polymer models of loop extrusion with barrier elements, and review recent experimental studies that provide strong support for loop extrusion, focusing on perturbations to CTCF and cohesin assayed via Hi-C in interphase. Finally, we discuss a likely molecular mechanism of loop extrusion by SMC complexes.

---

In interphase, mammalian chromosomes are folded into a series of insulated regions, termed topologically-associating domains (*TADs*), often elaborated with *peaks* at their corners, *grids* of peaks within and between TADs, and enriched lines (or *tracks*) of contact frequency emanating from a boundary (Fig. 1C, for review (Merkenschlager and Nora 2016; Bonev and Cavalli 2016)). TADs (Dixon et al. 2012; Nora et al. 2012), peaks (Rao et al. 2014), and tracks (Fudenberg et al. 2016) have an independent mechanistic origin from patterns associated with compartmental segregation of active and inactive chromatin (Schwarzer et al. 2017), and we discuss the interplay of these two mechanisms elsewhere (Nuebler et al. 2017). TAD boundaries are frequently demarcated by binding sites of the transcription factor ***CTCF,*** and are enriched for the Structural Maintenance of Chromosomes (SMC) complex ***cohesin.*** Functionally, TADs are believed to demarcate coherent *cis* neighborhoods of gene-regulatory activity and hence are crucial for development (Spielmann and Mundlos 2016). To explain how such neighborhoods can be formed, we put forward a mechanism based on a still-hypothetical process of ***loop extrusion.***

Here we present emerging evidence that interphase chromosomes are organized by loop extrusion, an active ATP-dependent process that allows nanometer-size molecular machines to organize chromosomes at much larger scales. We review how loop extrusion by cohesins can explain the formation of TADs, peaks, and tracks visible in interphase Hi-C maps. We then detail specific predictions made by the polymer model of loop extrusion, and discuss recent experimental perturbations to CTCF and cohesin that test these predictions and provide strong support to the loop extrusion mechanism. While we focus on comparisons to mammalian interphase Hi-C experiments, loop extrusion likely plays important roles in other organisms and parts of the cell cycle. We also discuss imaging experiments, single-molecule experiments, and a possible molecular mechanism of loop extrusion.

## Polymer model of loop extrusion with barrier elements

We frame our discussion around how we originally implemented the mechanism of loop extrusion limited by barriers as a polymer model (Fudenberg et al. 2016). In the process of loop extrusion, loop extruding factors ***(LEFs)*** translocate along the chromosomes, holding together progressively more genomically distant loci along a chromosome, thus producing dynamically expanding chromatin loops. LEF translocation is either halted by encounters with other LEFs, or probabilistically halted at specific genomic loci that contain ***extrusion barriers.*** We assume that if halted only on one side, a LEF may continue to extrude chromatin from its other side. LEFs continue to extrude until they dissociate from the chromatin fiber, releasing the extruded loop.

The minimal system of LEFs limited by extrusion barriers that we implement is defined by four parameters (Fig. 1A):

- ***lifetime*** on chromatin (sec)
- ***velocity*** along the chromatin fiber (kb/sec)
- ***separation*** between LEFs (kb)
- ***permeability*** of the extrusion barriers (probability)

**Figure 1.**
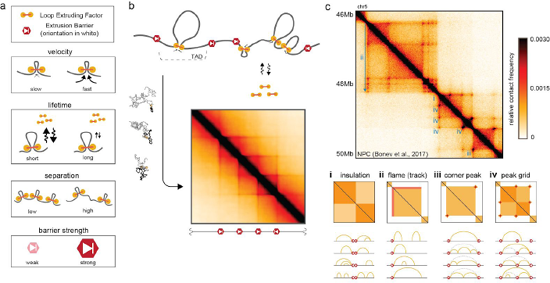
Polymer model of loop extrusion with barrier elements recapitulates features of interphase chromosome folding. a. Illustrations of the four key parameters governing the dynamics of interphase loop extrusion with barrier elements: LEF velocity, LEF lifetime, LEF separation, and barrier strength. Characterizing how changes to these parameters affect Hi-C maps ***in silico*** allows us to make experimental predictions for perturbations.
b. To compare our models with Hi-C experiments, we generate ensembles of conformations for each set of parameters, and then compute average contact maps. To compare with imaging experiments we can calculate other observables from the conformational ensemble.
c. Interphase Hi-C data from mouse neural progenitor cells (Bonev et al. 2017), annotated with features that can emerge via loop extrusion with barrier elements in blue ***(i-iv).*** Arc diagrams depict how stochastic configurations of LEF-mediated loops in distinct nuclei can lead to the population-averaged features. Chromatin loops directly held by LEFs are depicted with yellow arcs, while dashed grey arcs depict 'effective loops' from sets of adjacent LEFs. HiGlass views (Kerpedjiev et al. 2017): http://mirnylab.mit.edu/proiects/emerging-evidence-for-loop-extrusion.

i. Insulation, observed as squares along the diagonal of Hi-C maps (i.e. TADs), arises when extrusion barriers halt LEF translocation. LEFs then facilitate additional contacts within TADs, but not between TADs.
ii. Flames (or tracks), observed as straight lines often emerging from the borders of TADs, arise naturally in the loop extrusion model when LEFs become halted on one side at a barrier locus, while continuing to extrude from the other side (termed “lines” in (Fudenberg et al. 2016)).
iii. Peaks of enriched contact frequency often appear at the corners of TADs, and also often coincide with intersection points of flames. These peaks emerge as a result of LEFs being halted on both sides by extrusion barriers.
iv. Peak grids can emerge either when internal boundaries are skipped, or via transitive sets of LEF-mediated loops.

For comparison to ensemble-averaged Hi-C experiments, that capture a snapshot of contacts occurring at a particular point in time, it is also useful to define the product of lifetime and velocity, ***processivity*** (kb), which indicates the average size of a loop that a LEF would extrude if left unobstructed. Motivated by observations of CTCF motif orientations at TAD boundaries and at peaks (Vietri Rudan et al. 2015; Rao et al. 2014), we implement barriers as being ***directional,*** i.e. halting LEFs approaching it from only one side. Barriers can be modeled as either halting LEFs as long as the blocking factor is present, or stalling them until LEF dissociation. In our models, the permeability can be thought to represent the probability that a barrier locus is occupied by a blocking factor.

To compare predictions from our simulations with experiments, we generate a simulated ensemble of chromatin conformations for a given set of parameters (Fig. 1B). To accurately simulate features of chromatin folding at the scale of TADs we typically use monomers representing several nucleosomes to simulate 10-50Mb of chromatin (i.e. many times a typical TAD size). From these conformations we can extract experimentally-relevant observables (Imakaev et al. 2015). These include maps of contact frequency that can be compared to Hi-C contact maps, as well as distributions of spatial distances between pairs of loci, that can be compared with FISH experiments (Fudenberg and Imakaev 2017). From the simulated contact maps, we can then extract TADs, peaks, and contact frequency decay curves, *P(s)*, as done for experimental Hi-C maps. By comparing simulated and experimental features, we can then define a set of wild-type parameters, from which perturbations, and hence predictions, can be made.

The mechanism of loop extrusion limited by barriers recapitulates many features of interphase chromosome folding visible in Hi-C maps (Fig. 1C), including:

- TADs, regions of enriched contact frequency between neighboring barriers
- Tracks, lines emerging from one side of a barrier
- Peaks and grids of peaks, occurring between proximal barriers in *cis* but not between chromosomes
- Inwards-oriented CTCF motifs at TAD boundaries and at peak bases Further support comes from site-specific disruptions of TAD boundaries and peak bases, which respectively result in merging of adjacent TADs (Nora et al. 2012; Narendra et al. 2015; Rodríguez-Carballo et al. 2017) and orientation-dependent losses of peaks (de Wit et al. 2015; Guo et al. 2015; Sanborn et al. 2015). To our knowledge, no alternative mechanism of interphase chromosome organization currently agrees with all the above.

While we focus here on interphase loop extrusion, we note that loop extrusion by SMCs appears to have important consequences in mitosis (Naumova et al. 2013; Goloborodko et al. 2016a; Gibcus et al. 2018), where the term was coined and first mathematically modeled (Alipour and Marko 2012). The closely related concepts of reeling (Riggs 1990), facilitated tracking (Blackwood and Kadonaga 1998), loop expansion (Kimura et al. 1999) and progressive loop enlargement (Nasmyth 2001) have a rich history. Loop extrusion also appears relevant in bacteria (Wang et al. 2017; Gruber 2014; Wang et al. 2015). There are also related proposals for interphase loop extrusion (Sanborn et al. 2015; Nichols and Corces 2015; Brackley et al. 2018; Yamamoto and Schiessel 2017), which we discuss briefly below.

While the terms “contact”, “loop”, and “interaction” are often used interchangeably in the chromosome organization literature, they are often used to describe very different features of Hi-C contact maps (Forcato et al. 2017). To avoid ambiguity in the context of loop extrusion, we reserve the term “loop” in the narrow sense, for two regions of a continuous chromatin fiber brought together by a LEF at a given point in time. Indeed, simulations (Fudenberg et al. 2016; Doyle et al. 2014; Hofmann and Heermann 2015; Benedetti et al. 2014) and data analyses (Fudenberg and Imakaev 2017; Finn et al. 2017; Cattoni et al. 2017; Giorgetti et al. 2014) demonstrate that peaks in interphase Hi-C maps are not consistent with stable chromatin loops. Therefore, we refrain from using “loop” to describe any feature of Hi-C contact maps, as they likely emerge from sets of dynamically extruded loops that vary stochastically from cell-to-cell (**Fig. 1C, i-iv**).

### Challenges for testing models of loop extrusion

The stochastic nature of loop extrusion poses an experimental challenge for testing predictions from the model. Extruded loops are not directly visible via population-average Hi-C approaches because they are located at different genomic positions in different cells at any given time. Even with single-cell Hi-C methods an individual pair of loci linked by an extruding loop would not appear particularly different from any other captured contact. Visualization of extruded loops by microscopy is similarly challenging due to their continually changing locations both along the genome and in 3D space. Direct confirmation that a particular chromatin loop has been extruded ***in vivo*** will require methods that can simultaneously track multiple DNA loci as well as the loop extruders themselves. Nevertheless, much of the strongest evidence to date supporting the role of loop extrusion in interphase comes from changes in Hi-C maps upon perturbations that affect specific components of the loop extrusion machinery.

## Predictions from the model of interphase loop extrusion

To make experimental predictions, we must first identify components of the interphase loop extrusion machinery with their biological candidates. Several lines of evidence make us hypothesize that cohesin complexes play the role of LEFs, and CTCF plays the role of an extrusion barrier (Fudenberg et al. 2016). Cohesin is highly homologous to condensins, the main complexes responsible for compacting mitotic chromosomes, and is enriched at TAD boundaries in interphase. CTCF is enriched at TAD boundaries at preferentially-oriented motifs, and, compared with other transcriptional regulators, binds relatively stably to its cognate sites (for review (Hansen et al. 2018)). With these identities, we discuss how our model of loop extrusion predicts different outcomes for three perturbations: depletion of CTCF, depletion of cohesins, and increased processivity of cohesins (Fig. 2).

**Figure 2.**
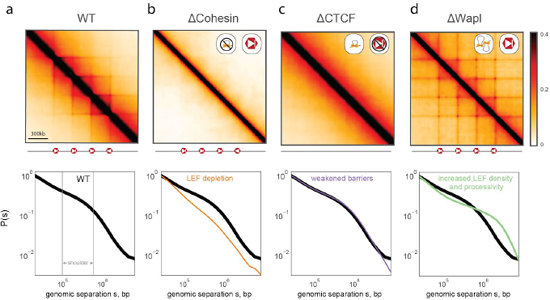
Loop extrusion polymer simulations predict the consequences of cohesin and CTCF perturbations. *Top row:* simulated Hi-C maps for indicated perturbations. ***Bottom row: P(s)*** for indicated perturbation compared to WT ***P(s).*** All simulations considered a 36Mb chain (3600 monomers) with the same positions and orientations of CTCF barriers (separated by 300kb) and the same LEF velocity (250 3D-per-1D steps). **a.**WT simulations used processivity 200kb, separation 200 kb, and barrier strength 0.995. The shoulder in ***P(s),*** indicative of comparction via loop extrusion, is indicated in grey. **b.**For ∆ Cohesin, our simulations predict the loss of TADs, peaks, flames, and the shoulder of ***P(s).*** ∆ Cohesin was simulated using: processivity 200kb, separation 2Mb, and boundary strength 0.995. This can represent the loss of actively extruding cohesins via ∆Nipbl, ∆Rad21, or other cohesin subunits. **c.**For ∆ CTCF, our simulations predict the loss of TADs, peaks, flames, yet no discernible change to ***P(s).*** This arises because CTCF plays an instructive role for the activity of extrusion. ∆ CTCF was simulated using processivity 200kb, separation 200kb, and boundary strength 0.9. **d.**For ∆Wapl, our simulations predict the emergence of additional peaks, including at further genomic separations, as well as an extension of the shoulder in ***P(s). ∆*** Wapl was simulated using processivity 1Mb, separation 150kb, and boundary strength 0.995.

### LEF depletion

For the depletion of the loop extruding factor, cohesin, our simulations display two phenomena (Fig. 2B) (i) the loss of TADs and associated Hi-C peaks; and (ii) decompaction of chromatin at the scales of individual loops (<200Kb). Changes in local compaction, in turn, can be studied by observing changes in the contact probability, ***P(s),*** as a function of genomic separation, ***s.*** Local compaction is seen as a region of ***P(s)*** with a shallow slope (~100-500kb), which we refer to as the ***shoulder*** (Fig. 2A); decompaction leads to reduction or loss of the shoulder region. We note that our models predict that a sharp decrease in LEF processivity would similarly lead to a loss of TADs, peaks, and compaction.

### Extrusion barrier depletion

For the depletion of site-specific extrusion barriers, as imposed by CTCF, our simulations also predict the loss of TADs and associated Hi-C peaks (Fig. 2C). However, our simulations predict other consequences of this perturbation should be very different from depletion of LEFs. This is because in our model, extrusion barriers only impose an instructive function on LEF translocation and positioning. We therefore predict little effect on overall compaction, and hence little change in the ***P(s)*** curve. This differentiates our predictions for CTCF depletion from those for cohesin depletion.

### Increased LEF density and processivity

For the depletion of a cohesin unloading factor, like Wapl, our model predicts that the consequent increased processivity and number of LEFs would lead to several phenotypes (Fig. 2D): (i) peaks at corners of TADs become stronger and appear between more distal barrier loci, creating extended grids of peaks; (ii) the orientational preference of barrier loci will become weaker, as LEFs halted at a directional barrier for long durations can stop traffic from the opposing direction as well. Finally, (iii), our model predicts that sufficiently increased loop extrusion will over-compact chromosomes. In Hi-C this would be detected as an extension of the shoulder in ***P(s),*** as opposed to how it recedes in the case of cohesin depletion. Macroscopically, sufficient compaction results in condensation into a prophase-like state, resulting in a cohesin-rich central scaffold for each chromosome.

Crucially, our model predicts that the loss of cohesin loop extruders and the loss of CTCF extrusion barriers should both lead to the loss of TADs and Hi-C peaks, yet in completely distinct fashions. Furthermore, increased processivity of cohesin extruders is predicted to manifest distinct phenotypes on Hi-C maps and macroscopic chromosome organization.

## Experimental perturbations consistent with interphase loop extrusion

While perturbing CTCF and cohesin dynamics is crucial for testing predictions of loop extrusion, depletion of such essential complexes poses many experimental challenges. For CTCF, cells begin dying after about 4 days of stringent depletion (Nora et al. 2017). For cohesin, there are additional challenges related to its essential role in sister chromatid cohesion and chromosome segregation during mitosis (Peters and Nishiyama 2012), and its multiple dynamically-exchanging subunits and regulators (Rhodes et al. 2017; Peters and Nishiyama 2012) that can be present in different abundances and likely have unique impacts on loop extrusion dynamics. Despite these challenges, recent studies have achieved modulation of cohesin and CTCF that result in dramatic changes, consistent with predictions from polymer models of loop extrusion (Table 1).

**Table 1:**
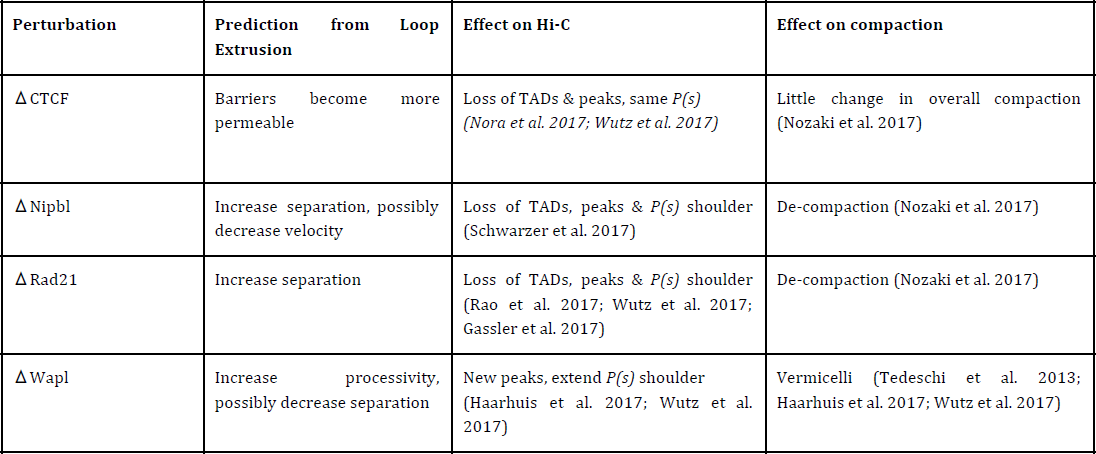
list of recent experimental perturbations, prediction from loop extrusion, effects in recent Hi-C experiments, and effect on overall chromatin density.

### Cohesin depletion

Consistent with our predictions for decreasing the number of active LEFs, depletion of the cohesin loader Nipbl (Scc2) (Schwarzer et al. 2017) and acute degradation of the cohesin kleisin Rad21 (Scc1) (Wutz et al. 2017; Rao et al. 2017) during interphase led to both: (i) complete erasure of TADs and Hi-C peaks (ii) and decompaction, as evidenced by loss of the ***P(s)*** shoulder (Fig. 3A). Decompaction is further supported by imaging, showing a reduction in H2B clustering by PALM following both RNAi knockdown of NIPBL and AID-mediated degradation of Rad21 (Nozaki et al. 2017). We note that earlier Hi-C studies (Sofueva et al. 2013; Zuin et al. 2014; Seitan et al. 2013) saw limited impact following the depletion of Rad21, potentially due to incomplete depletion.

**Figure 3.**
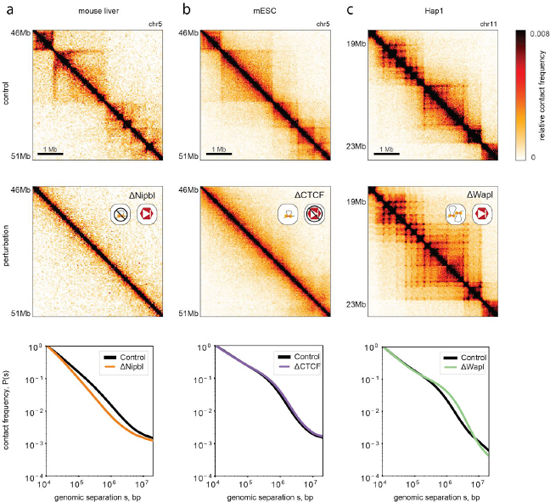
**Experimental phenotypes are consistent with predictions from loop extrusion simulations** *Top row:* unperturbed experimental Hi-C maps, replotted from indicated studies (see ***Methods***, for interactive HiGlass displays see: http://mimvlab.mit.edu/proiects/emerging-evidence-for-loop-extrusion.). *Middle Row:* Hi-C maps for indicated perturbations. *Bottom row: P(s)* for indicated perturbation compared to unperturbed ***P(s)*** normalized to contact frequency at 10kb. **a.**Schwarzer et al. used tissue-specific CRE-inducible gene deletion in mouse liver cells to deplete Nipbl (Schwarzer et al. 2017). **b.**Nora et al. used an auxin-inducible degron system to deplete CTCF in mESCs (Nora et al. 2017). **c.**Haarhuis et al. deleted Wapl in the Hap1 haploid human cell line, via CRISPR (Haarhuis et al. 2017).

A corollary of the Nipbl depletion result is that cohesin must be constantly loaded on chromatin to maintain TADs and associated corner-peaks. Consistently, TADs and Hi-C peaks are both rapidly lost upon AID-mediated degradation of Rad21 (<3hrs (Wutz et al. 2017)) and re-established after auxin washoff (40-60 minutes (Rao et al. 2017)). These consequences follow directly from our loop extrusion models, and the turnover time of cohesin (~5-30 min (Gerlich et al. 2006; Hansen et al. 2017; Wutz et al. 2017)).

Future studies, including a time course of Nipbl degradation, will be useful to dissect the dynamics of the processes and the potential role of Nipbl beyond that of a loader (Rhodes et al. 2017; Petela et al. 2017). In particular, while Nipbl depletion appears to have a dramatic effect on extrusion, knockout of its co-factor Mau2 (Scc4) appears to have a much weaker effect on loading yet a fairly strong effect on processivity (Haarhuis et al. 2017). Moreover, we note that different components of the interphase extrusion machinery could be limiting at different concentrations and in different contexts. We hypothesize that, via its consequences on loop extrusion, modulation of the levels of various cohesin subunits and interactors can serve to fine-tune overall gene regulation across cell-types and tissues.

### CTCF depletion

Consistent with our predictions for the loss of site-specific barriers to extrusion, acute auxin-induced degradation of CTCF in mESCs (Nora et al. 2017) and HeLa cells (Wutz et al. 2017) led to a dramatic loss of TADs and Hi-C peaks (Fig. 3B). However, the ***P(s)*** curve did not change, implying that while demarcation of contact-insulating boundaries in Hi-C maps was lost, the same degree of chromatin compaction was maintained. In support of the dynamic exchange of LEFs in our model, the effect of CTCF depletion was fully reversible following a two-day washoff period (Nora et al. 2017). We note that stringent dosage depletion was necessary to observe dramatic insulation defects: even a 15% preservation of CTCF showed a relatively mild phenotype (Nora et al. 2017). Similar loss of TADs and peaks were reported for ***in vivo*** inducible CTCF knockout in cardiomyocytes (Lee et al. 2017). Weaker effects have also been reported recently (Kubo et al. 2017; Rosa-Garrido et al. 2017) and earlier (Zuin et al. 2014), but this may have been due to relatively inefficient depletion or lower starting levels of CTCF.

The predicted lack of decondensation following CTCF depletion is further supported by imaging. PALM shows little difference in H2B clustering (Nozaki et al. 2017). Imaging of FISH probes at selected loci upon CTCF degradation show that inter-TAD distances increased while intra-TAD distances remained the same (Nora et al. 2017). Together these results are consistent with global compaction levels being unchanged but with diminished insulation across CTCF sites. Importantly, the lack of chromatin decompaction in CTCF depletion rules out models in which CTCF is strictly required for the loading (Nichols and Corces 2015) of chromatin-bound cohesin and any ensuing cohesin-mediated loops. Instead, the differences in imaging and Hi-C maps upon CTCF versus cohesin depletion are consistent with the loop extrusion model we describe (Fudenberg et al. 2016), where CTCF barriers serve an instructive function (Wendt and Peters 2009), and cohesin is loaded onto chromatin and can compact chromosomes through extrusion even in the absence of CTCF.

### Wapl Depletion

Consistent with our predictions for increasing the processivity and density of active LEFs, depletion of the cohesin unloader Wapl led to multiple phenotypes observed in Hi-C maps (Haarhuis et al. 2017; Wutz et al. 2017; Gassler et al. 2017) and by imaging (Tedeschi et al. 2013). For Hi-C (Fig. 3C) this includes (i) strengthened peaks at TAD corners, (ii) emergence of new peaks between boundaries at greater separations, creating extended grids of corner-peaks; (iii) a weakened correspondence between these features and CTCF motif orientation. Increased local compaction upon Wapl depletion is reflected by (iv) extension of the shoulder in the P(s) curve and, (v) the emergence of prophase-like vermicelli chromatids via imaging (Tedeschi et al. 2013). This remarkable observation provides further evidence for a ***universal molecular mechanism*** ‑‑ loop extrusion‑‑underlying both metaphase and interphase chromosome organization (Imakaev et al. 2015; Dekker and Mirny 2016).

Depletion of another component of the cohesin unloading machinery, Pds5A and Pds5B (Pds5A/B), led to many of the same phenotypes (Wutz et al. 2017), consistent with the increased the amount and residence time of chromatin-bound cohesin upon its depletion. However, there were also intriguing differences which may prove instructive for determining exactly how CTCF halts the progression of cohesin along the chromosome, e.g. Pds5 may instruct directional cohesin stalling (Petela et al. 2017; Wutz et al. 2017), and competition between the two HAWK family proteins, Nipbl and Pds5, may regulate cohesin translocation velocity (Petela et al. 2017). The observation that Wapl depletion appears to largely rescue the Hi-C phenotype of Mau2 depletion provides further support to the proposal that the Nipbl/Mau2 ‘loading complex' also has roles in promoting cohesin processivity for loop extrusion (Haarhuis et al. 2017). Finally, consistent with loop extrusion simulations with increased processivity, the joint depletion of Wapl and Pds5A/B showed even stronger effects in terms of shifting the shoulder in *P(s)* and in the emergence of vermicelli.

Collectively, the congruence of both Hi-C and imaging experiments following the perturbation of CTCF and cohesin dynamics strongly supports the role of loop extrusion in interphase. Future simulations and experiments will be valuable for probing the consequences of multiple simultaneous perturbations (Busslinger et al. 2017; Wutz et al. 2017).

## Single-molecule experiments support active loop extrusion

While providing strong support for chromosome folding by loop extension *in vivo*, the studies discussed above do not directly probe the molecular details of loop extrusion. Molecularly realizing the process of loop extrusion presents a considerable challenge, namely that the protein complexes performing loop extrusion need to ***track consistently*** in *cis* along chromatin, over large distances (up to tens-of-thousands of nucleosomes) without falling off. Moreover, the substrate, chromatin, is highly disordered due to nucleosomes and other DNA-bound proteins, likely posing a greater challenge than tracking along microtubules performed by cytoplasmic motors. Here we discuss how recent single molecule experiments argue that loop extrusion likely occurs via an active process, driven by molecular motors. While many of these observations were made with condensin and bacterial SMCs, they illustrate that loop-extrusion is a plausible mechanism of action for the whole family of SMC proteins, including cohesin.

### ATP-dependent Translocation

Recently, (Terakawa et al. 2017) demonstrated that a single yeast condensin has motor activity and is able to translocate processively along naked DNA *in vitro*. Using a DNA curtain assay, they found individual condensin complexes travel unidirectionally, rapidly (~4kb/min) and processively (~10kb) in an ATP-dependent manner with 10nm steps (30bp on naked DNA). As previous single-molecule studies only reported sliding dynamics of SMCs ((Stigler et al. 2016; Davidson et al. 2016; Kanke et al. 2016; Kim and Loparo 2016), for review (Eeftens and Dekker 2017)), the directional translocation observed by Terekawa et al. is incredibly important.

The high structural homology of cohesin to condensin makes it likely that the same physical mechanism would govern its processive motion, in addition to its established role of mediating sister chromatin cohesion (Peters and Nishiyama 2012). Indeed, the ability of these SMCs to compact chromosomes appears to be remarkably coherent over evolutionary timescales and cellular contexts (Schalbetter et al. 2017). Due to its dual roles, and more elaborate set of subunits, however, reconstituting this activity for cohesin may be more difficult *in vitro*. Nevertheless, we believe that the *in vitro* observations of ATP-driven processive condensin translocation argue against the likelihood of motor-free mechanisms (Brackley et al. 2018; Yamamoto and Schiessel 2017) of SMC processivity in general, including for cohesin.

While strongly supporting the loop extrusion mechanism, the single-molecule experiments leave open several questions of how loop extrusion can work *in vivo*:

- How can SMCs translocate on chromatinized rather than naked DNA?
- How can translocation result in loop extrusion?
- Is the measured speed of translocation sufficient to generate TADs and peaks?
- Do cells have sufficient ATP budgets to support extrusion during interphase?

### Walking hypothesis

In particular, it remains to be understood how SMC complexes can translocate on chromatin fibers rather than naked DNA. Translocations while maintaining constant contact with DNA may not always be possible due to the complexity of chromatin fiber and abundance of DNA-bound proteins. Although the size of an SMC complex (~50nm) exceeds that of a single nucleosome (~10nm), nucleosomes would constitute challenging obstacles for SMC translocation if maintaining constant contact with DNA is required for translocation.

A possible solution comes from the structural similarity of SMC domain organization to that of kinesin and myosin motors (Peterson 1994; Guacci et al. 1993) that walk on microtubules and actin, which suggests that SMCs can similarly walk on chromatinized DNA. Importantly, a ***walking mechanism*** would allow translocation where obstacles such as nucleosomes and other DNA-bound proteins can be passed over, avoiding disruptions of the underlying nucleosomal array. During each step of the walking process, one SMC head can remain DNA-bound, while the other hops forward and rebinds nearby DNA (Fig. 4A). SMC walking is consistent with the rapid and flexible dynamics of their arms (Eeftens et al. 2017), and the 10nm step size (Terakawa et al. 2017) would allow passing over nucleosomes (e.g. by hopping from linker to linker) and other DNA-bound complexes, avoiding the need for unwinding nucleosomal DNA or nucleosome eviction (Fig. 4B).

**Figure 4.**
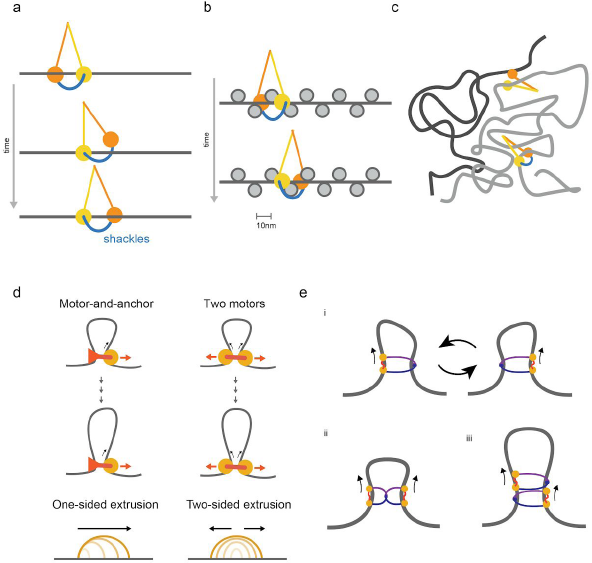
**a.**Walking as a possible mechanism of SMC translocation, with SMC arms in yellow and orange, kleisin in blue, creating a ***shackled walker.*** **b.**Walking along a chromatin fiber, by hopping from linker-to-linker without disrupting nucleosomal DNA. **c.**Benefit of topological entrapment: an SMC walker without a kleisin can step from one chromatin strand (grey) to another in its vicinity (black), whereas a shackled SMC walker with a kleisin is able to track in ***cis*** over long distances. **d.**Two possible mechanisms for converting translocation to extrusion; the first involves a single translocating motor attached to an anchor, leading to single-sided extrusion. The second involves two motors translocating in opposite directions, leading to two-sided extrusion. **e.**Possible realizations of motor activity by SMCs ***(i-iii). i.*** a single SMC acting as single motor that switches between entrapped chromatin strands, effectively performing two-sided extrusion; ***ii.*** dimerized SMCs performing two-sided extrusion; ***iii.*** alternatively dimerized SMCs performing two-sided extrusion.

A walking mechanism would be greatly aided by the known ability of SMCs to ***topologically entrap*** DNA (Peters and Nishiyama 2012), which can ensure that the walker tracks in ***cis,*** along the same chromatin fiber (Fig. 4C). Pseudo-topological (Srinivasan et al. 2017) entrapment can similarly help maintain extrusive cohesins on the same DNA molecule over long genomic distances (Kschonsak et al. 2017). In other words, SMC complexes may translocate along the chromatin fiber as ***shackled walkers*** (Fig. 4A). Below we discuss how translocation DNA can result in loop extrusion.

An important open question is how CTCF, and possibly other chromatin-bound proteins, can halt cohesin translocation while nucleosomes do not, when they are fairly similar in size. While they probed diffusive sliding rather than processive tracking dynamics, (Davidson et al. 2016) report that cohesin can rapidly slide over some DNA-bound proteins and nucleosomes, but becomes obstructed by DNA-bound CTCF and transcriptional machinery; a similar, yet more restrictive, dependence of sliding on the size of DNA-bound factors has been reported in other single-molecule studies probing sliding dynamics (Davidson et al. 2016; Kanke et al. 2016). This suggests that CTCF blocks translocation of cohesin by a specific mechanism rather than by steric exclusion, e.g. by inhibiting the ATPase action of the cohesin machinery (Petela et al. 2017; Wutz et al. 2017) directly or via other cohesin interactors (e.g. via Pds5) and potentially in concert with co-factors (Hsu et al. 2017). Alternatively, CTCF may recruit additional co-factors to increase its physical size or pose a greater challenge for walking due to its DNA binding geometry (Hashimoto et al. 2017).

### From translocation to loop extrusion

Multiple possibilities exist as to how the translocation of motor complexes along a chromosome can realize the process of loop extrusion. These include: (1) a single translocating motor attached to a chromatin anchor; (2) two connected motors translocating in opposite directions; (3) a single motor that switches between two chromatin fiber substrates (Fig. 4D). These architectures for the action of SMC motors can lead to different consequences for the processive dynamics of extrusion. Uni-directional extrusion could result from a single motor-and-anchor architecture. Bi-directional extrusion would emerge from the latter two possibilities. We note there are multiple possibilities for how many SMC complexes are required to realize motor activity (Fig. 4E), either as monomers or, potentially, oligomers (Keenholtz et al. 2017). One advantage of two-motor extrusion is that it naturally allows one motor to continue extruding if the other becomes blocked. Since models discussed here and in (Fudenberg et al. 2016; Sanborn et al. 2015; Goloborodko et al. 2016a, 2016b) assumed independent bidirectional extrusion, it remains unclear if one-sided loop extrusion is sufficient to form TADs, peaks, and tracks, as well as for compacting mitotic chromatids.

### Velocity of loop extruders

The measured rates of stepping and step sizes for condensin (Terakawa et al. 2017) agree well with the expectations of the loop extrusion theory in interphase for cohesin. Using ~2 steps per second and ~10nm step size measured ***in vitro,*** this gives ~18kb/min if cohesin moves one nucleosome per step (~150bp). This is further doubled if cohesin extrusion occurs via a two-motor mechanism, yielding ~36kb/min. These values are compatible with the ~10-30kb/min predicted by polymer models as sufficient to generate TADs and corner peaks ***in vivo.*** There are several ways to arrive to this estimate. The first involves dividing the size of the largest TADs (~1Mb, (Bonev and Cavalli 2016)) by time to re-establish of TADs following exit from mitosis (~0.5-2h (Naumova et al. 2013), ~30min (Nagano et al. 2017)) or following auxin wash-off (~30min (Rao et al. 2017)). Alternatively, one can use the processivity of cohesin estimated from fitting Hi-C data with loop extrusion models (~200-400Kb (Fudenberg et al. 2016)), and divide this by the cohesin turnover time (~5-30min (Gerlich et al. 2006; Hansen et al. 2017; Wutz et al. 2017)).

We note that pushing by RNA Pol II alone, at its reported velocities, would be too slow (~1.5-3kb/min (Veloso et al. 2014; Jonkers et al. 2014; Danko et al. 2013)). The observation of cohesin-dependent features in both active and inactive chromatin (Schwarzer et al. 2017; Haarhuis et al. 2017), as well as the transcriptionally inactive maternal zygotic pronucleus (Gassler et al. 2017), further argues against Pol II providing the primary motive force for loop extrusion.

### Energy Budget

A simple estimate shows that the energy burden of ATP consumption by loop-extruding cohesins in interphase is negligible as compared to ATP production in a mammalian cell. Again using 2 ATP per sec per SMC complex (Terakawa et al. 2017), and the total number of actively extruding cohesin molecules, either measured (~30,000-100,000 per cell (Hansen A.S., personal communication)) or estimated from fitting simulations to Hi-C data (~1 loop-extruder per 200Kb, i.e. ~60,000 per diploid G2 cell), one obtains a very low rate of ATP consumption (<210^5^ ATP per sec). This constitutes less than 0.02% of the 10^9^ ATP/sec production rate by a fibroblast (Flamholz et al. 2014). Thus the energy burden of chromosome organization by cohesin is marginal.

## Conclusions

While the key role of molecular motors in the cytoplasm is broadly appreciated (Phillips et al. 2012), there is now a growing appreciation for loop extrusion by SMC complexes as an active processes organizing and compacting chromatin in the nucleus (Nasmyth 2017; Haarhuis and Rowland 2017). Analogous to the myriad uses for the contractile dynamics of active actin and tubulin networks, we hypothesize that the interphase loop extrusion has been repurposed for a variety of biological ends (Fudenberg et al. 2016; Dekker and Mirny 2016), including targeting VDJ recombination, and regulation of enhancer-promoter interactions.

### Additional interactive media available at

http://mirnylab.mit.edu/proiects/emerging-evidence-for-loop-extrusion.

## Acknowledgements

The authors thank Elphege Nora and Anders Stejr Hansen for detailed feedback. This work was supported by NIH (GM114190) and NSF, Physics of Living Systems (15049420) grants and by the Center for 3D Structure and Physics of the Genome of NIH 4DN Consortium (DK107980). GF was supported by the San Simeon Fund (PI: K. Pollard).

## Methods

### Hi-C analysis

Published Hi-C datasets were re-processed using the *Distiller* workflow https://github.com/mirnylab/distiller-nf), producing filtered pairs files https://github.com/4dn-dcic/pairix/blob/master/pairsformatspecification.md) and *Cooler* contact matrix files https://github.com/mirnylab/cooler. (Abdennur et al. 2017)). *P(s)* curves were calculated on Hi-C pairs using logarithmically increasing genomic distance bins. For display, filtered bins were imputed via nearest-neighbor interpolation. Interactive *HiGlass* (Kerpedjiev et al. 2017) displays for relevant datasets are provided at http://mirnylab.mit.edu/proiects/emerging-evidence-for-loop-extrusion.

### Simulations of loop extrusion with extrusion barriers

Loop extrusion with barrier element dynamics were modeled as described previously (Fudenberg et al. 2016), using an updated translocator (https://bitbucket.org/mirnylab/openmm-polymer/src/8534bc3183e0727a83cdcb9b5525736774035884/examples/loopExtrusion/smcTranslocator.pyx. described in (Fudenberg and Imakaev 2017)). Polymer dynamics were simulated using OpenMM (Eastman et al. 2013; Eastman and Pande 2010), as described previously (Fudenberg et al. 2016). All simulations considered a 36Mb chain (3600 monomers) with the same positions and orientations of CTCF barriers (separated by 300kb) and the same LEF velocity (250 3D-per-1D steps). WT simulations used processivity 200kb, separation 200 kb, and pausing barrier strength 0.995. To simulate perturbations, simulations were run with modified parameters. ΔCohesin was simulated using: processivity 200kb, separation 2Mb, and boundary strength 0.995. ΔCTCF was simulated using processivity 200kb, separation 200kb, and boundary strength 0.9. ΔWapl was simulated using processivity 1Mb, separation 150kb, and boundary strength 0.995.

**Table S1.** list of exper mental perturbation, likely consequences for loop extrusion, cell type and reference where Hi-C for this perturbation was performed.

| Perturbation | Putative function | Study | Perturbation method | Cell type | Reported effects |
| --- | --- | --- | --- | --- | --- |
| $\Delta$ Rad21 | extruder subunit | Sofueva et al., 2013 | conditional Cre-mediated deletion (Cre-ERT2 or transfected Cre) | NSCs and ASTs derived from mouse ESC line | Mild effect on Hi-C ma, including relaxed domain structure" |
|  |  | Seitan et al., 2013 | conditional Cre-mediated deletion (CD4-Cre) | Differentiating mouse T-cells | Mild effect on Hi-C map, and resilience of compartment assignments |
|  |  | Zuin et al., 2014 | HRV protease-induced cleavage | HEK239T human hepatocyte cell line | Mild effect on Hi-C map, though with reduced interactions within TADs |
|  |  | Gassler et al., 2017 | conditional Cre-mediated deletion (Zp3-Cre) | Fertilized mouse oocytes | Loss of aggregate TADs and peaks. Loss of P(s) shoulder |
|  |  | Rao et al., 2017 | auxin-inducible degron | HCT-116 human colorectal carcinoma | Loss of TADs and peaks. |
|  |  | Wutz et al., 2017 | auxin-inducible degron | HeLa | Loss of TADs, peaks and P(s) shoulder |
| $\Delta$ Nipbl | loader | Schwarzer et al., 2017 | conditional tissue-specific Cre-mediated deletion (Ttr-Cre/Esr1) | mouse liver | Loss of TADs, peaks, and P(s) shoulder. |
| $\Delta$ Mau2 | loading cofactor | Haarhuis et al., 2017 | CRISPR-mediated knockout | HAP1 haploid human cell line | Weakening and shortening of detectable TADs and peaks. P(s) shoulder recedes. |
| $\Delta$ CTCF | barrier | Zuin et al., 2014 | RNAi | HEK239T human hepatocyte cell line | Mild effect on Hi-C map, though with an increase in interactions between neighboring domains |
|  |  | Nora et al., 2017 | auxin-inducible degron | E14TG2a mouse ESC line | Loss of TADs and peaks, same P(s) |
|  |  | Kubo et al., 2017 (bioRxiv) | auxin-inducible degron | F123 mouse ESC line | Weakened TADs, reduced interaction frequency at peaks |
|  |  | Lee et al., 2017 (bioRxiv) | conditional tissue-specific Cre-mediated deletion (transfected cTnt-Cre) | mouse heart | Loss of TADs & peaks |
|  |  | Rosa-Garrido et al., 2017 | conditional tissue-specific Cre-mediated deletion (MerCreMer) | mouse heart | Loss of some peaks, maintenance of TADs |
|  |  | Wutz et al., 2017 | auxin inducible degron | HeLa | Loss of TADs & peaks, same P(s) |
| $\Delta$ Wapl | unloader | Haarhuis et al., 2017 | CRISPR-mediated knockout | HAP1 haploid human cell line | Increased extent of TADs and peak grids, stronger peaks, P(s) shoulder shifts to the right. |
|  |  | Gassler et al., 2017 | conditional Cre-mediated deletion (Zp3-Cre) | Fertilized mouse oocytes | Stronger aggregate TADs and peaks. P(s) shoulder shifts to the right |
|  |  | Wutz et al., 2017 | RNAi | HeLa | Increased extent of TADs, peak grids, stronger peaks, P(s) shoulder shifts to the right. |
| $\Delta$ Pds5A/B | unloading cofactor | Wutz et al., 2017 | RNAi | HeLa | Increased extent of TADs, peak grids, stronger peaks, P(s) shoulder shifts to the right. |

